# The marine gastropod *Crepidula fornicata* remains resilient to ocean acidification across two life history stages

**DOI:** 10.1101/2020.10.08.331967

**Authors:** Christopher L Reyes, Brooke E Benson, Morgan Levy, Xuqing Chen, Anthony Pires, Jan A Pechenik, Sarah W Davies

## Abstract

Rising atmospheric CO_2_ reduces seawater pH causing ocean acidification (OA). Understanding how resilient marine organisms respond to OA may help predict how community dynamics will shift as CO_2_ continues rising. The common slipper shell snail *Crepidula fornicata* is a resilient marine gastropod native to eastern North America, which has been a successful invader along the western European coastline and elsewhere. To examine its potential resilience to OA, we conducted two controlled laboratory experiments. First, we examined several phenotypes and genome-wide gene expression of *C. fornicata* in response to pH treatments (7.5, 7.6, 8.0) throughout the larval stage and then tested how conditions experienced as larvae influenced juvenile stages (i.e. carryover effects). Second, we examined genome-wide gene expression patterns of *C. fornicata* larvae in response to acute (4, 10, 24 and 48 hours) pH treatment (7.5, 8.0). Both *C. fornicata* larvae and juveniles exhibited resilience to OA and gene expression responses highlight the role of transcriptome plasticity in OA resilience. Larvae did not exhibit reduced growth under OA until they were at least 4 days old. These phenotypic effects were preceded by broad transcriptomic changes, which likely serve as an acclimation mechanism for combating reduced pH conditions frequently experienced in littoral zones. Delayed metamorphosis was observed for larvae reared at reduced pH. Although juvenile size reflected larval rearing pH conditions, no carryover effects in juvenile growth rates were observed. Transcriptomic analyses suggest increased metabolism under OA, which may indicate compensation in reduced pH environments. Time course transcriptomic analyses suggest energetic burdens experienced under OA eventually dissipate, allowing *C. fornicata* to reduce metabolic demands and acclimate to reduced pH. This study highlights the importance of assessing the effects of OA across life history stages and demonstrates how transcriptomic plasticity can allow highly resilient organisms, like *C. fornicata*, acclimate to reduced pH environments.

## Introduction

Rising atmospheric carbon dioxide (CO_2_) concentrations, resulting from anthropogenic emissions, are causing substantial increases in the acidity of the world’s oceans (Bigg, Jickells, Liss, & Osborn, 2003; IPCC, 2013). This process, known as ocean acidification (OA), involves carbonic acid formed by the hydrolysis of atmospheric CO_2_ in seawater dissociating into bicarbonate [HCO_3_^-^] and hydrogen ions [H^+^], lowering seawater pH (Orr et al., 2005). Average ocean pH has declined by 0.13 units since 1765 and is expected to decrease a further 0.3-0.4 units by 2100, corresponding to an atmospheric CO_2_ concentration of 800-1000 ppm (IPCC, 2001; Orr et al., 2005). Impacts on the phenology, recruitment, and community interactions of organisms across diverse geographical regions are expected to strengthen as climatic conditions continue to shift (Walther et al., 2002). Additionally, nearshore ecosystems (e.g. estuaries and intertidal zones) characterized by substantial daily fluctuations in salinity, temperature, and pH are likely to experience even greater impacts in the future (Baumann, Wallace, Tagliaferri, & Gobler, 2015; Diederich & Pechenik, 2013; Pacella, Brown, Waldbusser, Labiosa, & Hales, 2018; Waldbusser & Salisbury, 2014).

A number of negative impacts of OA on the physiology of marine mollusks have been well documented (Dineshram et al., 2015; Maboloc & Chan, 2017; Vargas et al., 2013; Waldbusser et al., 2016). As acidity increases, excess H^+^ recombines with carbonate to form bicarbonate. This reduction in the availability of carbonate ions, which are required by mollusks and other calcifying organisms for shell formation, has consequences for growth, morphology, and ultimately survival (Orr et al., 2005; Talmage & Gobler, 2010). Additionally, it can reduce fertilization success, compromise induced defenses, and impair immune function (Vargas et al., 2013). Although most species exhibit reduced fitness under elevated CO_2_ concentrations (Guo, Huang, Pu, You, & Ke, 2015; Talmage & Gobler, 2010), sensitivity to acidification can vary between related species and even among different populations of the same species (Griffiths, Pan, & Kelly, 2019; Guo et al., 2015; Noisette, Bordeyne, Davoult, & Martin, 2016; Talmage & Gobler, 2010). The molecular basis for these patterns remain poorly characterized.

Native to the eastern coast of North America, the common slipper shell snail *Crepidula fornicata* has demonstrated resilience to most environmental stressors associated with climate change (Blanchard, 1997; Kriefall, Pechenik, Pires, & Davies, 2018; Noisette et al., 2016). Adults of this species have been shown to tolerate elevated *p*CO_2_ (750 μatm) for >5 months and exhibit decreased calcification rates only under the extreme *p*CO_2_ of 1400 μatm (Noisette et al., 2016). However, the early life history stages may be more vulnerable to elevated CO_2_ (Guo et al., 2015; Talmage & Gobler, 2010). Noisette et al. (2014) showed that *C. fornicata* larvae reared at reduced pH exhibited reduced shell growth and reduced mineralization and also showed signs of shell abnormalities. In addition, while there was no consistent influence of pH on larval mortality within the range of pH 7.5-8.0, reduced pH negatively impacted larval and juvenile growth rates and delayed the onset of competence for metamorphosis (Bogan, McMahon, Pechenik, & Pires, 2019; Kriefall et al., 2018; Pechenik et al., 2019). Short- and long-term exposure of competent larvae to reduced pH (7.5 vs. 8.0), however, did not inhibit metamorphosis in response to a natural adult-derived cue (Pechenik et al., 2019). The physiological mechanisms underlying such resistance are not well understood.

Although these studies have shed light on how this resilient mollusk copes with OA, few studies have investigated how the impacts of OA may interact across life stages in marine mollusks, despite the complexities associated with life stage transitions (Bogan et al., 2019; Ross, Parker, & Byrne, 2016; Talmage & Gobler, 2011). Here, we investigate how exposure to OA in the larval stage may influence later life stages (i.e., latent effects or carryover effects) by assessing the physiological and transcriptomic responses of *C. fornicata* larvae reared at three pH levels (pH 7.5, 7.6, 8.0). After metamorphosis, juveniles were raised under control conditions (pH 8.0), and the influence of pH conditions experienced as larvae on juvenile growth and patterns of gene expression were examined. In addition, we determined how quickly these patterns of gene expression were affected by acute pH exposure, by comparing transcriptomic responses of larvae reared at pH 8.0 (control) and pH 7.5 after 4 hours (h), 10 h, 24 h, and 48 h. We demonstrate dynamic responses across life stages, highlighting *C. fornicata*’s resilience and capacity for acclimatization to drastically altered environmental conditions.

## Materials and Methods

### Adult and larval collection

Brooding adults of *C. fornicata* were collected during their reproductive season in June and July 2017 from the intertidal zone in Totten Inlet, Thurston Co., WA and transported to the University of Washington’s Friday Harbor Laboratories (FHL) in Friday Harbor, WA. Stacks of approximately four to six adults were housed in separate, aerated 3-L glass jars containing 2 L of room temperature (~ 23 °C) unfiltered seawater, which was changed daily. Larvae hatched naturally within several days after adults were collected. Veligers were concentrated by gently siphoning the culture through a 150 μm sieve shortly after their release by the mothers. Veligers from each jar were released by a different female, and thus were considered to be separate broods. Each of the two experiments described in detail below was conducted independently to test different questions.

### Seawater pH manipulation and carbonate chemistry

Larvae were cultured in the Ocean Acidification Environmental Laboratory at FHL. Incoming seawater was filtered to 1 μm and equilibrated overnight at 20°C by bubbling with mixtures of CO_2_ and CO_2_-free air delivered by Aalborg GFC17 mass-flow controllers to achieve pH levels of 7.5, 7.6, and 8.0 (Kriefall et al., 2018). Seawater pH was measured immediately before loading into culture jars with a Honeywell Durafet pH electrode calibrated to the total scale by the cresol purple spectrophotometric method described by Dickson, Sabine, and Christian (2007). Headspaces of culture jars were continuously ventilated with the same gas mixtures used to condition the seawater pH treatments. Temperature and salinity were measured with a YSI Pro Series 1030 meter. Seawater samples were fixed with mercuric chloride and titrated to determine total alkalinity (TA) using a Mettler DL15 automated titrator calibrated to certified reference materials (Dickson laboratory, Scripps Institution of Oceanography). *p*CO_2_ and Ωar were estimated based on empirical measurements of pH and TA using CO_2_Sys 2.1 (Pierrot, Lewis, & Wallace, 2006). Chemical and physical properties of larval culture seawater are given in Table S1 and Table S2.

### Experiment I: Larval culturing and growth measurements

150 larvae from a single brood were randomly assigned to each of four replicate 800-mL jars per pH treatment (7.5, 7.6, or 8.0) and reared under treatment pH conditions for 12 days (d). Every 2 d, larvae were isolated by sieving and then transferred back into jars with freshly conditioned seawater. Larvae were fed 10 × 10^4^ cells mL^-1^ of *Isochrysis galbana* (clone T-ISO) when cultures were started and at each water change, to promote maximal growth rates (Pechenik & Tyrell, 2015). A subsample of approximately 25 larvae was collected from each replicate jar on days 4 and 8 and stored in RNAlater (Thermo Fisher Scientific) for gene expression profiling.

Veliger larval growth rates were quantified according to three metrics: change in shell length (μm), proportion tissue weight relative to total weight, and change in total weight (ng). Random subsamples of 20 larvae from each experimental replicate (*n* = 80 larvae/treatment) were collected soon after hatching (day 0) and 4, 8, 11, and 12 days later to document shell length growth rates (μm/day). Larvae were nondestructively imaged using a Motic camera fitted to a Leica Wild M3C dissecting microscope and then returned to culture. Shell lengths were measured in ImageJ (Schneider, Rasband, & Eliceiri., 2012). Growth rates were recorded cumulatively relative to initial measurements from day of hatching (day 0). At 4- and 8-d, 20 additional larvae per replicate were fixed in 20% buffered formalin before being transferred into pre-weighed aluminum cups to assess changes in shell and tissue weight. Samples were oven dried at 60 °C for at least 6 h, weighed, and then placed into a muffler furnace at approximately 500 °C for 6 h to oxidize all organic material, allowing for determinations of larval tissue weights and inorganic shell weights.

### Induction and measurement of competence for metamorphosis

Larval competence for metamorphosis was assayed on days 10 and 12 to estimate the impact of reduced pH on time to competence before all larvae in all cultures were induced to metamorphose. In each experimental assay, 30 larvae per replicate (*n* = 120 larvae/treatment) were transferred to 8 mL of seawater that matched larval pH rearing conditions and were then exposed to 20 mM elevated KCl to induce metamorphosis (Pechenik & Gee, 1993). Metamorphosis was defined by the loss of the ciliated velum, and the proportion of metamorphosed larvae was assessed 6 h after induction.

### Juvenile culturing and growth measurements

Metamorphosed larvae (hereafter, “juveniles”) were maintained for an additional 4 d at ambient conditions (pH 7.9-8.0, salinity 29-30 ppt, temperature 22-23° C) on a diet of T-ISO at 18 x 10^4^ cells/ml. Groups of 5-20 juveniles were subsampled for gene expression immediately following metamorphosis on day 11 or 13, approximately 24 h (1 d) after their respective induction trials, as well as 4-d post-metamorphosis (DPM), at 14-d or 16-d. Collections were pooled across days such that all individuals at 1-DPM or 4-DPM were analyzed together, in order to standardize time spent in the juvenile stage at the time of sampling. All samples were preserved in RNAlater (Thermo Fisher Scientific). Mean juvenile shell length (μm) was quantified upon metamorphosis using Image J as described for larvae, and at 4-DPM using an ocular micrometer at 25x magnification. These data were then used to calculate juvenile shell length growth rates (%/day) between 1-DPM and 4-DPM.

### Experiment II: Larval culturing

An second study was conducted to determine how quickly larvae respond to reduced pH at the molecular level. Culturing methods for this short-term (48-h) experiment were similar to those for larvae (reared for 12-d) described above. Briefly, larvae were sourced from a separate brood and cultured for 2 d from hatching at pH 8.0 as described above (4 replicate cultures, 200 larvae/replicate). After 2 d, larvae from each culture were equally divided into 2 new culture jars containing freshly-conditioned seawater at pH 7.5 or pH 8.0. Larvae were fed as described above, subsampled (approximately 25 larvae from each of the 4 replicates of each pH treatment, *n* = approximately 100 larvae /treatment) after 4, 10, 24, and 48 h, and stored in RNAlater (Thermo Fisher Scientific) for gene expression profiling. No phenotypic measurements were taken for this secondary acute OA experiment.

### Statistical analyses of phenotypic effects

To determine if significant differences in larval growth rates (change in shell length and proportion tissue weight) and larval shell lengths were observed between treatments over the course of the experiment, a one-way analysis of variance (ANOVA) followed by a Tukey’s Honest Significant Difference (HSD) test was performed for each time point independently. One-way ANOVAs were also used to compare rates of metamorphosis across treatments, average juvenile shell lengths, and shell length growth rates between 1- and 4-DPM. Again, Tukey’s HSD tests were used to test for differences between levels within pH treatment. Assumptions for all parametric models (normality and equal residuals) were assessed via diagnostic plots. All data visualization and analyses were implemented in R (R Core Team, 2018).

### RNA isolation and sequencing preparation

In samples from the 16-d experiment, RNA was isolated from a pool of approximately 15 individuals per replicate jar (*n* = 60 individuals/treatment) for larval gene expression (4- and 8-d) and approximately 5 individuals per replicate jar (*n* = 20 individuals/treatment) for juveniles (1- and 4-DPM). Total RNA was extracted from all samples using RNAqueous Total RNA Isolation Kit (Invitrogen) per manufacturer’s instructions with the following modification: 0.5 mm glass beads (Sigma Aldrich, Z250465) were added to lysis buffer and samples were homogenized using a bead beater as per Kriefall et al. (2018). Trace DNA contamination was eliminated using *DNase1* (Invitrogen, AM2222) and gel electrophoresis confirmed RNA integrity and the absence of trace DNA. Approximately 500 ng of DNased total RNA was used to prepare tag-based RNAseq libraries, excluding samples that failed to yield at least 10 ng of DNased total RNA (*n* = 13; 9 larval, 4 juvenile). Libraries were prepared following Meyer, Aglyamova, & Matz (2011) with appropriate modifications for Illumina Hi-Seq sequencing (Dixon et al., 2015; Lohman, Weber, & Bolnick, 2016). Prepared libraries (*n* = 35) were sequenced across two lanes of Illumina Hi-Seq 2500 (50bp single-end dual-indexed reads) at the Tufts University Core Facility (TUCF). RNA was similarly isolated from larvae in the 48-h experiment. Replicates constituted a pool of approximately 15 individuals per jar (*n* = 60 individuals/treatment), and libraries (*n* = 36) were prepared, as described previously, from 500 ng DNased RNA and sequenced separately from the 16-d experiment across two Illumina Hi-Seq 2500 (50bp single-end dual-indexed reads) lanes.

### Transcriptome assembly and read mapping

Due to low mapping efficiencies to the previously used *Crepidula fornicata* transcriptome (Henry, Perry, Fukui, & Alvi, 2010; Kriefall et al., 2018), a novel *C. fornicata* transcriptome was assembled using all sequencing data from the current study. Data from all libraries were pooled, which yielded 533.8 million single-end reads. *Fastx_toolkit* was used to trim *Illumina TruSeq* adapters and poly(A)+ tails, and reads were then quality filtered using *fastq_quality_filter* with the requirement that ≥ 80% of bases meet a cutoff score of at least 20. These trimmed reads then served as input for RNAseq *de novo* assembly using Trinity (Grabherr et al., 2011) with 750 GB of system memory and 24 CPUs at the Shared Computing Cluster (SCC) at Boston University (BU). Contigs > 200bp in length were annotated by BLAST sequence homology searches against UniProt and Swiss-Prot NCBI NR protein databases with an e-value cutoff of e^-5^ and annotated sequences were subsequently assigned to Gene Ontology (GO) categories (The UniProt Consortium, 2015). The transcriptome and its associated annotation files can be accessed at http://sites.bu.edu/davieslab/files/2019/05/Crepidula_fornicata_transcriptome_2019.zip.

As with the transcriptome, *fastx_toolkit* was used to remove *Illumina TruSeq* adapters and poly(A)^+^ tails from each individual sequence library. Short sequences (<20bp) and low-quality sequences (quality score < 20) were also trimmed. A customized perl script was used to remove PCR duplicates sharing the same degenerate header and transcript sequence (https://github.com/z0on/tag-based_RNAseq). Resulting quality-filtered reads were then mapped to the newly assembled *C. fornicata* transcriptome using *Bowtie2.2.0* (Langmead & Salzberg, 2012) and a per-sample counts file was created using a custom perl script. The script summed up reads for all genes and discarded reads that mapped to multiple genes. Mapped reads for the 16-d experiment ranged from 330,871 to 1,557,855 with mapping efficiencies ranging from 40.5% to 45.0% (Table S3), and reads for the 48-h experiment ranged from 94,004 to 1,235,041 with mapping efficiencies ranging from 40.7% to 46.2% (Table S4). It should be noted that these values exclude samples with low read counts, which were removed during downstream outlier analyses.

### Gene expression analyses

Differential gene expression analyses were performed with *DESeq2* v. 1.22.2 (Love, Huber, & Anders, 2014) in R v. 3.5.1 (R Core Team, 2018). Data were tested for outliers using the package *arrayQualityMetrics* (Kauffmann, Gentleman, & Huber, 2009), and any samples failing two or more outlier detection methods (*n* = 5 for each experiment, Tables S3 & S4) were excluded from subsequent analyses and generally reflected those samples with low depth of coverage. Differentially expressed genes (DEGs, FDR: 0.05) between the reduced pH treatments (pH 7.5 and 7.6 in the 16-d experiment and pH 7.5 in the 48-h experiment) relative to the control (pH 8.0) were identified at each independent time point (4- and 8-d and 1- and 4-DPM in the 16-d experiment and 4-, 10-, 24-, and 48-h in the 48-h experiment) using generalized linear models (design= ~ treatment) and p-values for the significance of these contrasts were generated using Wald statistics. These p-values were then adjusted using the false discovery rate method (Benjamini & Hochberg, 1995).

Data from original per-sample counts files were normalized to total counts and log-transformed using the *decostand* function. These data were then used as input for canonical correspondence analyses with the package *vegan* (Oksanen, Blanchet, Kindt, Legendre, & O’Hara, 2016) to characterize overall differences in global gene expression across treatments. Overall significance of constraints (day/hour, treatment) was assessed with the *anova* function, which conducted a permutation test and constructed sampling distributions by resampling the data.

Gene ontology (GO) enrichment analyses were performed using Mann-Whitney U tests on ranked p-values (GO-MWU, Voolstra et al., 2011). GO enrichment was used to group together sets of genes as categories, based on their function, under the overarching divisions of ‘cellular component’ (CC), ‘biological process’ (BP), and ‘molecular function’ (MF). Enrichment analyses allowed for a general overview of which categories were being differentially regulated under the reduced pH conditions. Negative logged p-values, used as a continuous measure of significance, were ranked and significantly enriched GO categories were identified. Results were plotted as dendrograms with hierarchical clustering of GO categories based on shared genes. Fonts and text colors were used to distinguish significance and direction of enrichment (up or down) for the regulated categories relative to pH 8.0. Gene annotation information, GO designation, raw mapped data counts, and *DESeq2* and GO enrichment results for both experiments can be accessed as supplemental files at https://github.com/chrislreyes/Crepidula.

## Results

### Effects of reduced pH on larval growth, metamorphosis, and juvenile shell growth

Larvae reared at pH 7.5 and pH 7.6 grew significantly slower than those reared at pH 8.0 over the course of 12 days, although shell length growth rates were not significantly different for the first 8 d (Figure 1A). By day 11, larval shell lengths were at least 30% higher for larvae reared at the control pH than for those reared at either of the reduced pH levels (F_2,9_ = 21.15; *P* < 0.001; Figure 1A). While larval tissue weight as a proportion of total weight (tissue weight/total weight) did not differ significantly between treatments at day 4, larvae reared at pH 8.0 had a significantly higher tissue weight proportion by day 8 relative to larvae reared at pH 7.5 or 7.6 (F2,7 = 6.0; *P* = 0.0304; Figure 1B). Larval metamorphosis was significantly affected by rearing pH: a significantly smaller percentage of larvae reared at pH 7.5 and 7.6 metamorphosed during both 6-h induction periods (10- and 12-d) relative to larvae reared at pH 8.0 (*P* < 0.003; Figure 2).

**Figure 1.**
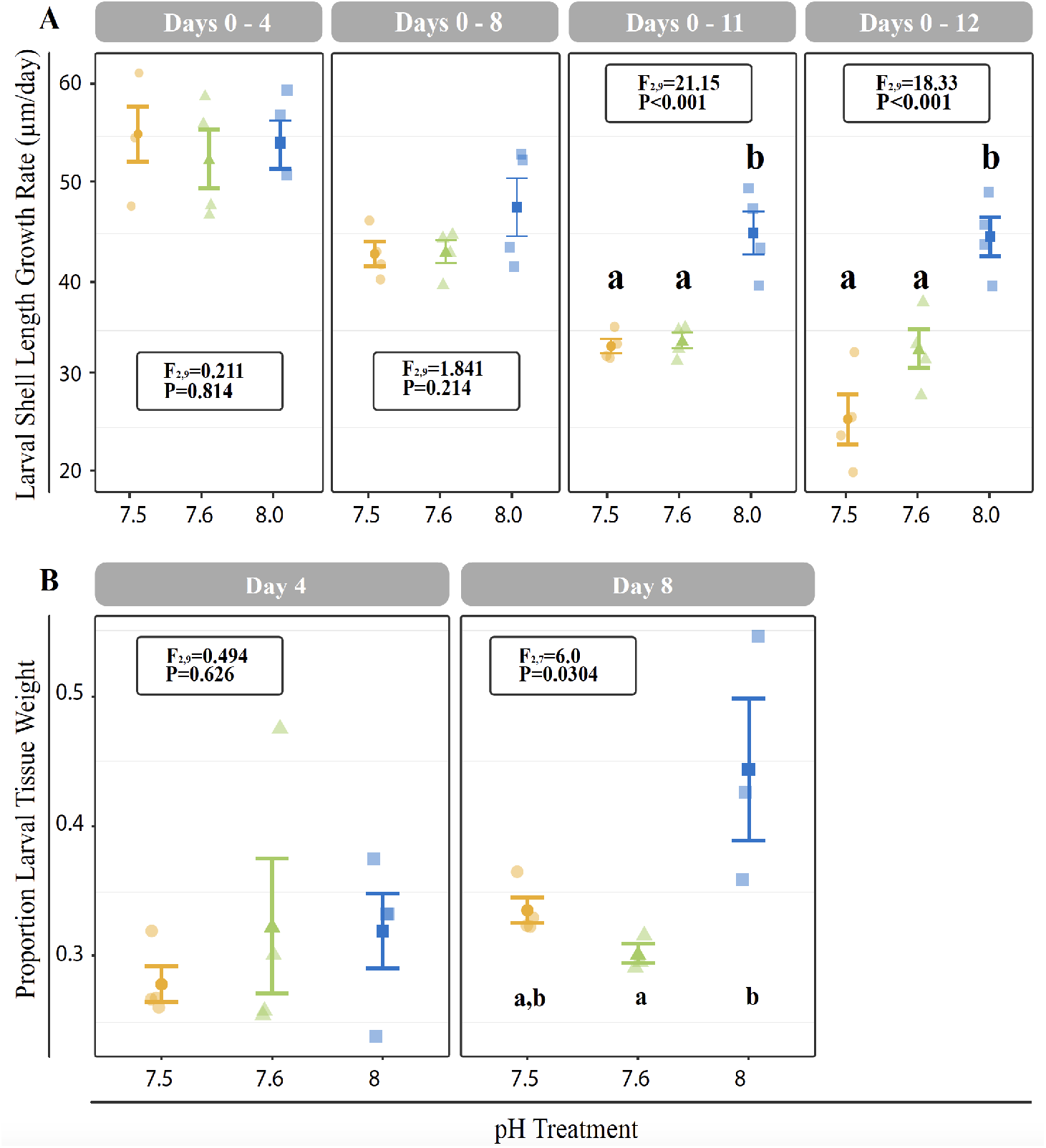
(A): Cumulative shell length growth rate (μm/day) for larvae relative to initial (day 0) measurements at 4-, 8-, 11-, and 12-d. (B): Proportion of tissue weight relative to total weight on days 4 and 8. Error bars represent +/- one standard error and different letters for pH treatments indicate significantly different means based on Tukey’s HSD tests (*P* < 0.05).

**Figure 2.**
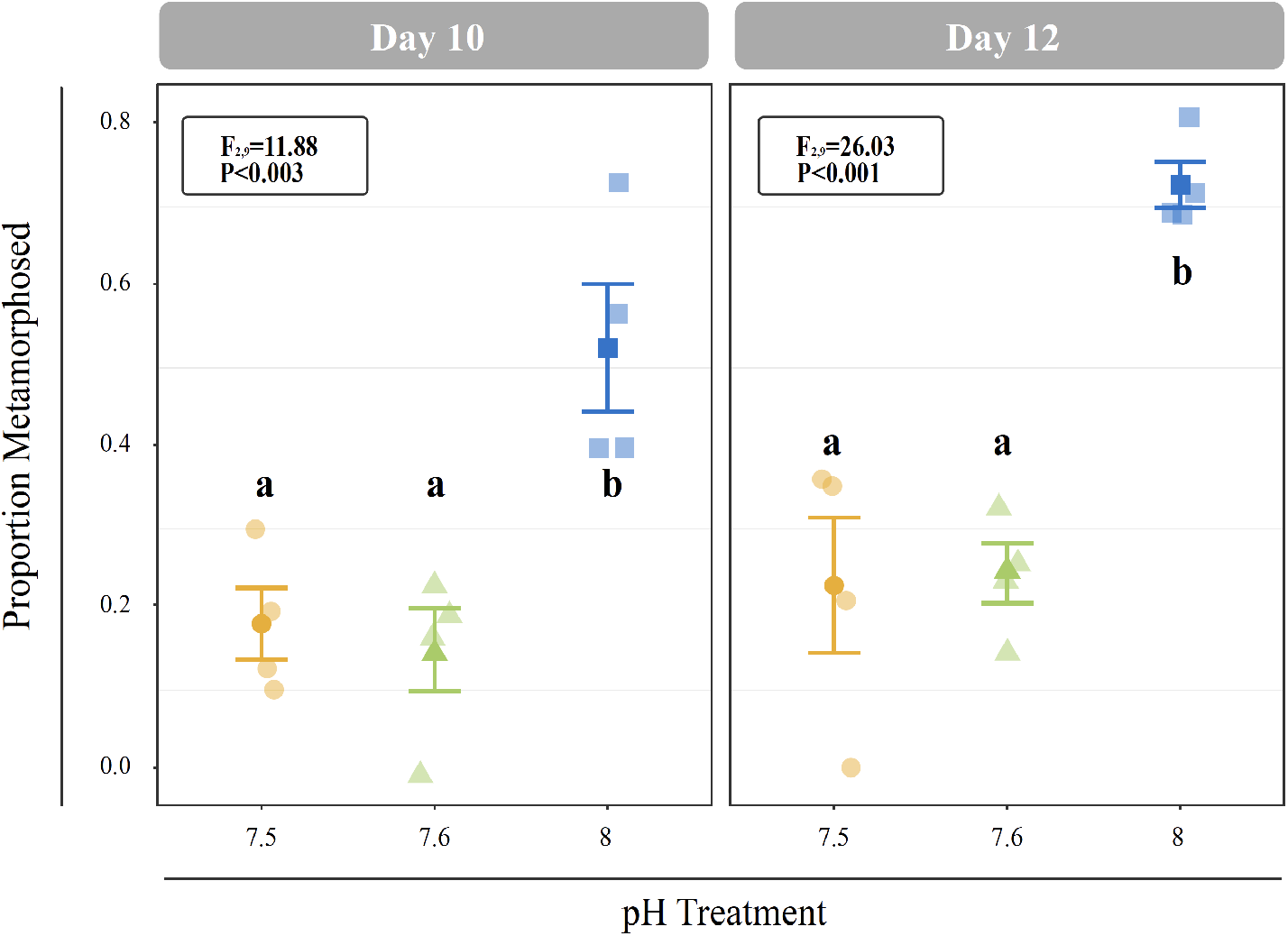
Proportion of larvae that metamorphosed after 6 h of induction with 20 mM elevated KCl at 10- and 11-d. Error bars represent +/- one standard error and different letters for pH treatments indicate significantly different means based on Tukey’s HSD tests (*P* < 0.05).

Juveniles reared at reduced pH as larvae had significantly smaller shell lengths at both 1-DPM and 4-DPM in comparison to juveniles reared at ambient pH as larvae (*P* < 0.001; Figure S1B-C). However, these reduced juvenile shell lengths were due to reduced larval shell lengths observed from day 8 onwards (Figure S1A) and significant differences in juvenile shell length growth rates were not observed after multiple test correction (*P*=0.0479; Tukey’s HSD >0.05; Figure 3A).

**Figure 3.**
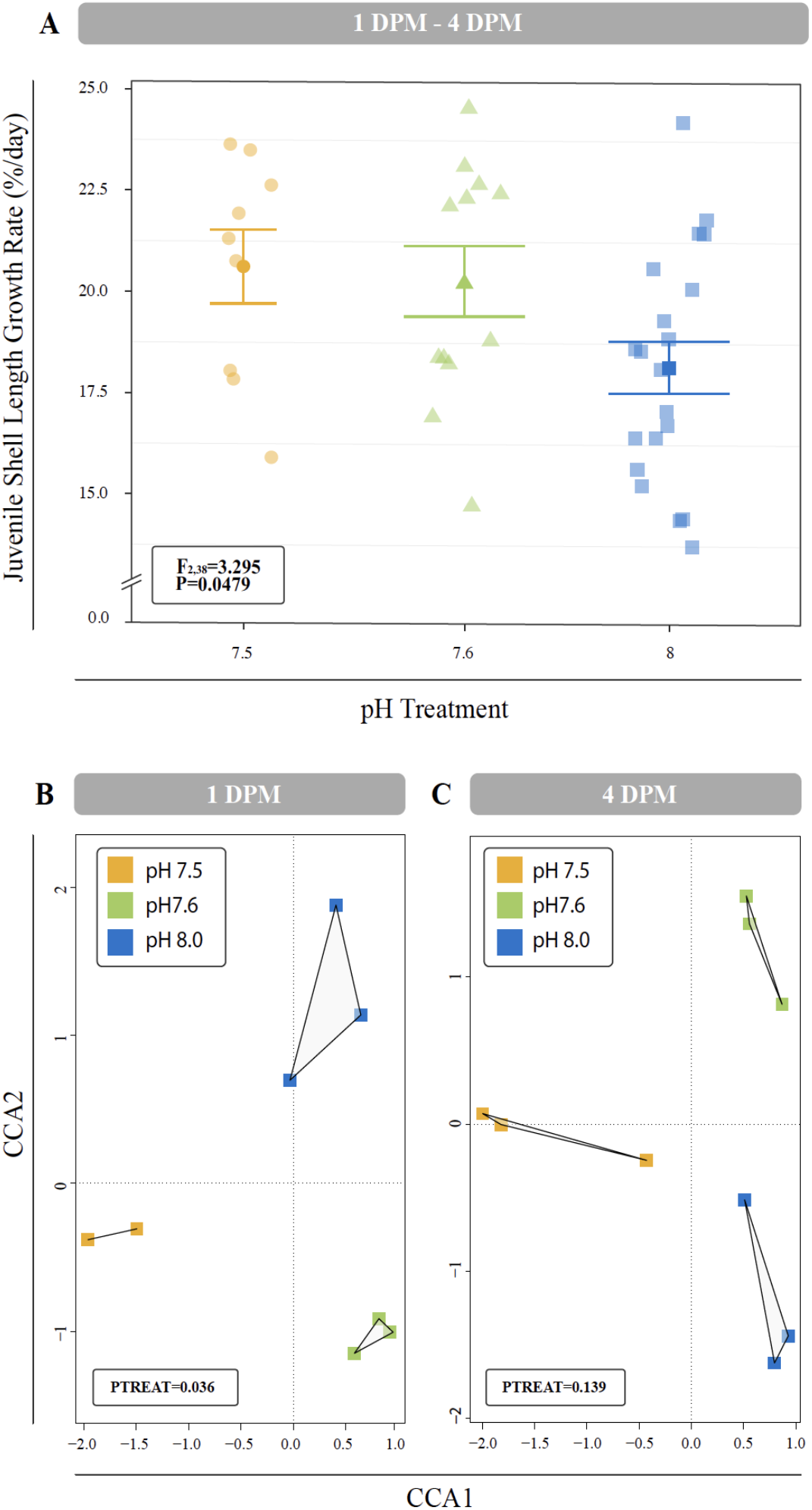
(A): Mean percent shell length growth rate (%/day) for juveniles between 1-DPM and 4-DPM. Error bars represent +/- one standard error. Overall difference between pH treatments was significant (*P* = 0.049), however no Tukey’s HSD test passed the 0.05 p-value cut-off suggesting no significant differences between any two levels of a factor. (B-C): Canonical Correspondence Analysis (CCA) of all log transformed isogroups clustered by experimental treatment at 1-DPM (B) and 4-DPM (C). Responses of C. *fornicata* juveniles across pH treatments was found to be significant at 1-DPM (*P* = 0.036), but not at 4-DPM (*P* = 0.139). Colors indicate pH treatment condition: blue = pH 8.0, green = pH 7.6, yellow = pH 7.5.

### Transcriptomic responses of Crepidula fornicata to reduced pH

Differences in global gene expression in *C. fornicata* larvae were better explained by larval age (4-d v. 8-d) than by pH treatment (Figure S2B), although neither of these factors resulted in significant sample clustering. In contrast, larval pH treatment had a greater influence on juvenile snail gene expression when compared to age, but again these differences were not significant (Figure S2C). Overall gene expression profiles did differ significantly between larval pH treatments in juveniles at 1-DPM (*P* = 0.036). These differences, however, were no longer significant by 4-DPM (*P* = 0.139; Figure 3B-C), suggesting *C. fornicata* acclimation. While there were trends in global gene expression differences between pH treatments, numbers of differentially expressed genes (DEGs) were generally very low, especially in larvae. Relative to larvae reared at pH 8.0, larvae reared at pH 7.5 only downregulated a single gene at 8-d and no DEGs were detected for larvae reared at pH 7.6 at 4- or 8-d. Juveniles that had been reared as larvae at pH 7.5 differentially expressed 92 genes (65 upregulated; 27 downregulated) at 1-DPM and 210 genes (140 upregulated; 70 downregulated) at 4-DPM in comparison to juveniles that had been reared as larvae at pH 8.0, and juveniles that had been reared as larvae at pH 7.6 differentially expressed 23 genes (2 upregulated; 21 downregulated) at 1-DPM but only a single gene (upregulated) at 4-DPM.

### GO enrichment in response to reduced pH of *Crepidula fornicata* larvae

In *C. fornicata* larvae, most GO enrichments were observed to be overrepresented under reduced pH treatments relative to pH 8.0 (red text), and there were consistently more GO enrichments detected in 4 day old larvae relative to 8 day old larvae (Figures 4 & S3). GO categories associated with ribosomal proteins (i.e. *ribosomal subunit;* GO:00443912, *small ribosomal subunit;* GO:00159354, *structural constituent of the ribosome;* GO:00037352) were consistently downregulated at 4- and 8-d in larvae reared under pH 7.5 in comparison to larvae reared at pH 8.0 (Figure 4) and in 4-d larvae reared under pH 7.6 (Figure S3). GO categories associated with mitochondrial proteins (i.e. *mitochondrial part*; GO:00444292, *respirasome*; GO:00704692, *mitochondrial membrane;* GO:0005743) were also found to be downregulated in 4-d larvae reared under pH 7.5 and in 4- and 8-d larvae reared under pH 7.6. In contrast, terms associated with cytoskeleton proteins (i.e. *cytoskeleton;* GO:00058562, *cytoskeleton organization;* GO:00070102, *regulation of cytoskeleton organization;* GO:00514932) were consistently upregulated in 4- and 8-d larvae reared under pH 7.5 and in 4-d larvae reared under pH 7.6. Additionally, GO categories associated with oxidative stress responses (i.e. *oxidoreductase activity*; GO:00164912, *oxidoreductase, acting on NAD(P)H;* GO:00166512, *oxidation-reduction process;* GO:00551142) were downregulated in 4-d larvae reared under pH 7.5 and in 4- and 8-d larvae reared under pH 7.6.

**Figure 4.**
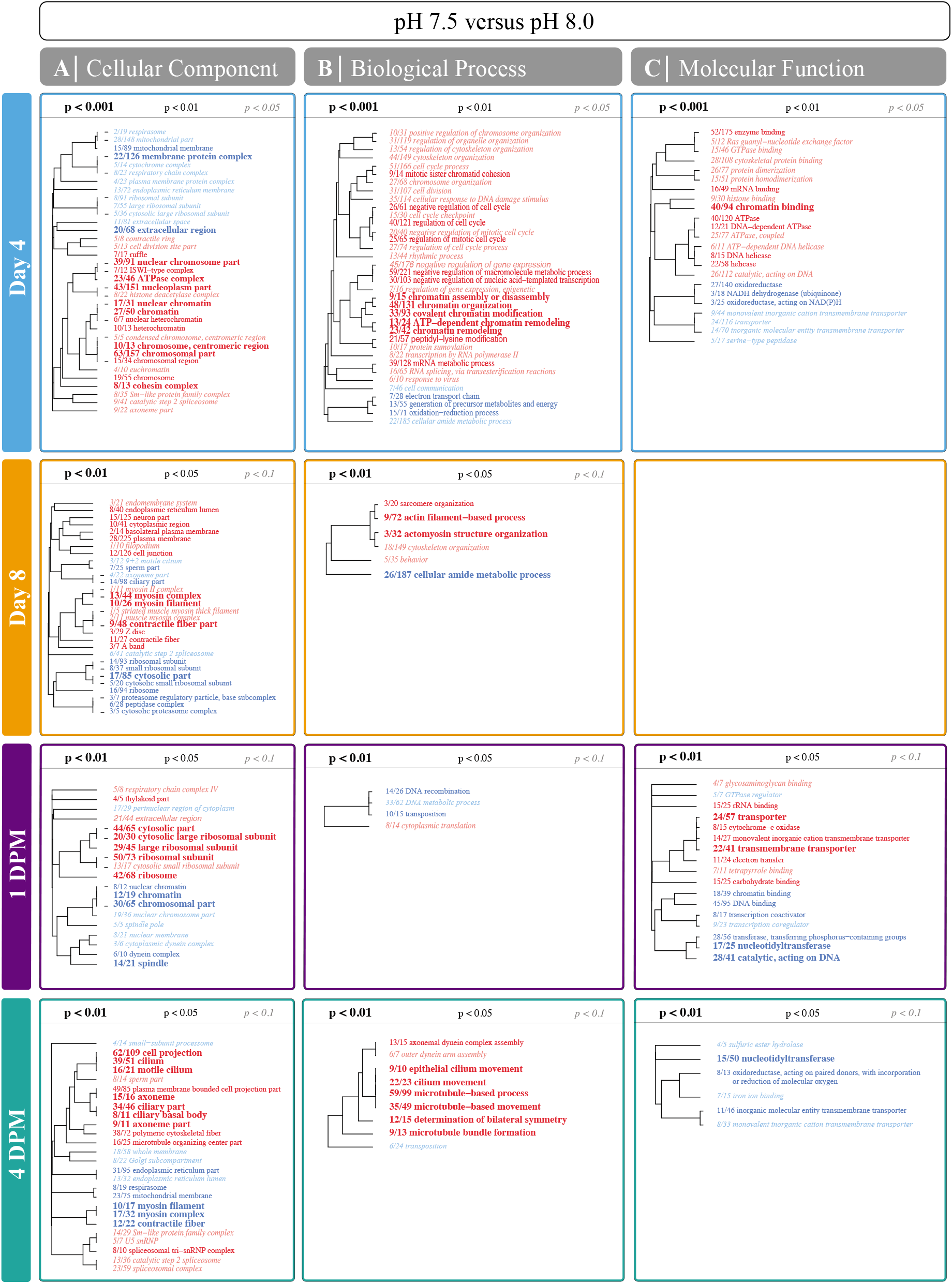
Significantly enriched gene ontology (GO) categories for the pairwise comparison between pH 7.5 and pH 8.0 treatments for larvae and juveniles in the 16-d experiment. Mann-Whitney U (MWU) tests were conducted based on ranking of signed log p-values and results were plotted as dendograms with an indication of genes shared between categories. Enrichment by ‘cellular component’, ‘biological process’, and ‘molecular function’ (columns) are shown for 4- and 8-d for larvae and for 1-DPM and 4-DPM (rows) for juveniles. Overrepresented categories relative to pH 8.0 are colored as red and underrepresented categories are colored as blue. A blank grid indicates that there were no significantly enriched categories for that division at that time point. Results for pH 7.6 and 8.0 can be found in supplemental Figure S3.

### *Crepidula fornicata* juvenile GO enrichment in response to larval rearing at reduced pH

In contrast to larval responses, most GO enrichments detected for juveniles derived from larvae reared at pH 7.6 were found to be largely underrepresented (blue text; Figure S3) while those derived from larvae reared at pH 7.5 were under- and overrepresented (Figure 4). Juveniles derived from larvae from both reduced pH treatments showed enrichment of ribosomal protein GO categories at 1-DPM, whereas those derived from larvae reared at pH 7.6 downregulated those same categories at 4-DPM (Figures 4 & S3). Similarly, juveniles derived from larvae reared at pH 7.6 downregulated categories associated with the cytoskeleton at 1-DPM (i.e. *polymeric cytoskeletal fiber;* GO:0005874, *microtubule-based process;* GO:00070172, *microtubule-based movement;* GO:00070182), yet juveniles derived from larvae reared at pH 7.5 exhibited enrichment of those same categories at 4-DPM (Figures 4 & S3). In contrast to larvae, juvenile *C. fornicata* derived from larvae reared at pH 7.5 upregulated mitochondrial proteins at 1-DPM. Juveniles derived from larvae reared at pH 7.6, however, downregulated these same GO terms at 4-DPM, again demonstrating the more dynamic responses of juveniles. Lastly, GO categories associated with oxidative stress responses were not differentially enriched at 1-DPM, but were downregulated at 4-DPM in juveniles derived from larvae reared at pH 7.5 only.

### 48-h reduced pH experiment: short term larval transcriptomic responses and GO enrichment

As observed for larvae in the 16-d experiment, differences in global gene expression for larvae in the 48-h experiment were better explained by larval age than by pH treatment (Figure S2A). Relative to pH 8.0, larvae reared at pH 7.5 differentially expressed 2 genes (1 upregulated; 1 downregulated) at 4 h and 33 genes (18 upregulated; 15 downregulated) at 48 h (FDR-adjusted < 0.1), while no genes were differentially expressed at 10 or 24 hours. In larvae reared under pH 7.5, GO categories associated with ribosomal proteins were underrepresented at 4 h, overrepresented at 10 h and 24, h and then underrepresented again at 48 h relative to larvae reared at pH 8.0 (Figure 5). GO categories associated with mitochondrial proteins were overrepresented at 4 h and 10 h and then underrepresented at 48 h in larvae reared under pH 7.5. Lastly, categories associated with oxidative stress responses were overrepresented at 10 h but these same categories were underrepresented at 48 h. Overall these responses suggest that larval *C. fornicata* exhibit initial transcriptomic responses to reduced pH, but these responses are reversed after 48 hours in treatment, demonstrating the transcriptome plasticity of larval *C. fornicata* in response to acute reduced pH.

**Figure 5.**
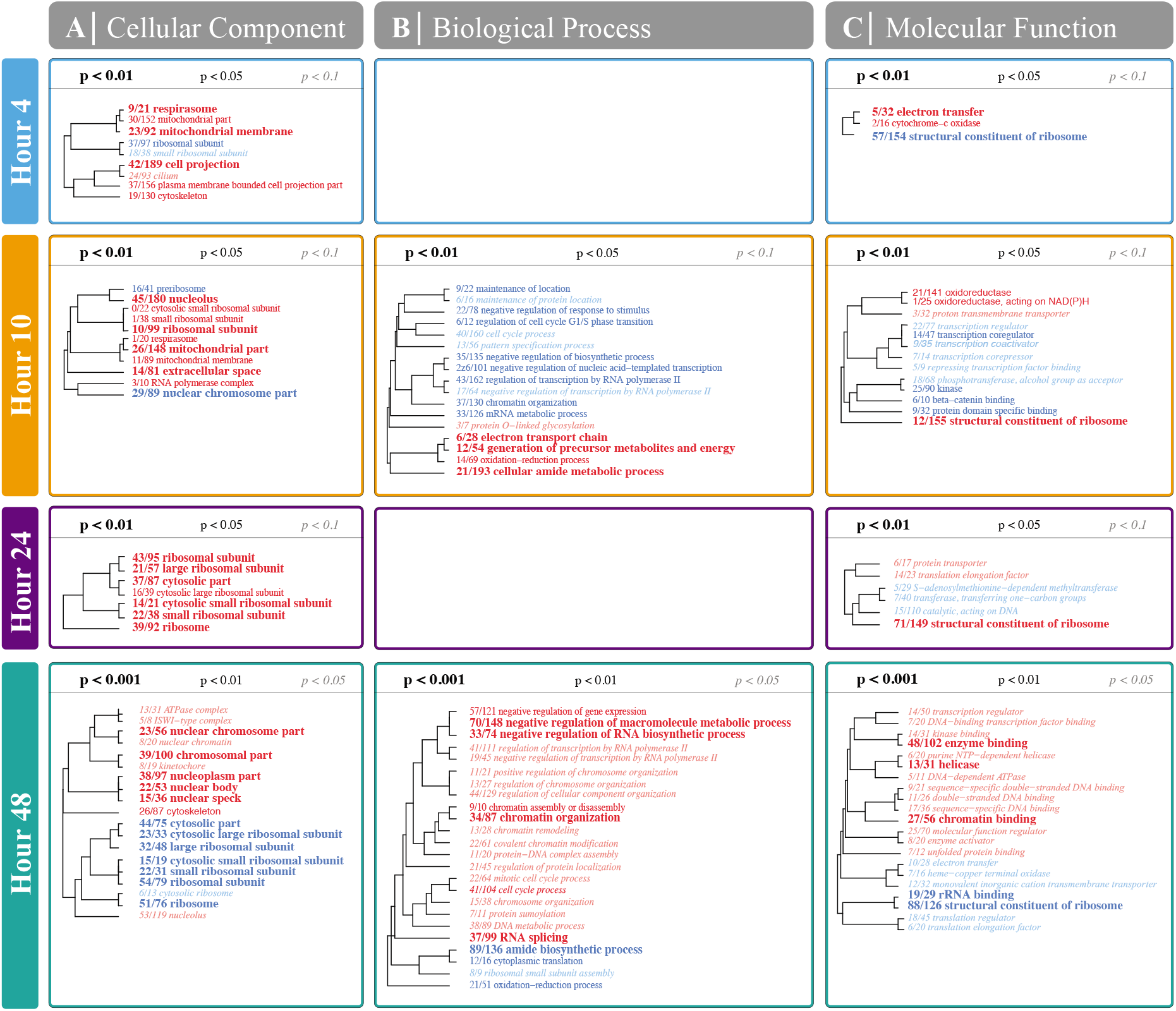
Significantly enriched gene ontology (GO) categories for the pairwise comparison between pH 7.5 and pH 8.0 treatments for the 48-h experiment larvae. Mann-Whitney U (MWU) tests were conducted based on ranking of signed log p-values and results were plotted as dendograms with an indication of genes shared between categories. Enrichment by ‘cellular component’, ‘biological process’, and ‘molecular function’ are shown for 4-, 10-, 24-, and 48-h. Overrepresented categories relative to pH 8.0 are colored as red and underrepresented categories are colored as blue. A blank grid indicates that there were no significantly enriched categories for that division at that hour.

## Discussion

### Reduced pH reduced larval growth in *Crepidula fornicata*

Consistent with other studies investigating the effects of reduced pH on larval growth rates in *C. fornicata* (Bogan et al., 2019; Kriefall et al., 2018), we observed reduced shell and tissue growth rates for larvae reared at reduced pH. These reductions were not immediately apparent and only became evident after 8 days of exposure for tissue growth and after 11 days of exposure for shell length increases (Figure 1). These results corroborate previous studies on *C. fornicata*’s congener *Crepidula onyx*, which first exhibited significantly slower larval growth after 14 days of rearing at reduced pH (7.3 and 7.7; Maboloc & Chan, 2017). Altered growth patterns in response to reduced pH have also been found in other organisms, including coral and algae, whose growth were similarly affected after two weeks of rearing at various elevated *p*CO_2_ treatments (55, 70, 100, and 210 Pa; Comeau, Edmunds, Spindel, & Carpenter, 2014). In contrast, no changes in shell length or development were found for embryos of the Olympia oyster *Ostrea lurida* after exposure to pH conditions as low as 7.4 (Waldbusser et al., 2016). It is worth noting, however, that these oyster embryos were only reared at reduced pH conditions for 5 days. In our study, 8 days of prolonged exposure to reduced pH was required before *C. fornicata* larvae exhibited detectable phenotypic responses. It is clear from this study and others that the amount of time spent under treatment (i.e. acute vs chronic) is a particularly important consideration when assessing an organism’s response to OA.

It has been previously suggested that reducing investment in calcification may minimize the energetic burden of living in reduced pH environments and serve as a potential mechanism of acclimation and resilience (Waldbusser et al. 2016). Indeed, the potential to acclimate to OA and maintain calcification rates has been proposed to be energetically costly (i.e. Cohen & McConnaughey, 2003). Previous work on larval sea urchins exposed to reduced pH, found reduced growth coupled with increased metabolic rates (Stumpp, Wren, Melzner, Thorndyke, and Dupont 2011). Davies, Marchetti, Ries, and Castillo (2016) also demonstrated that reduced calcification rates in adult corals were coupled with increased expression of genes associated with metabolism. Similarly, Comeau et al. (2014) noted that rapidly calcifying corals were more sensitive to the impacts of OA than slow calcifiers. Overall, our results suggest that larval *C. fornicata* may alter their growth after experiencing prolonged reduced pH in order to minimize their energetic burden, which may explain the exceptional resilience of this species in response to a variety of stressors (Diederich & Pechenik, 2013; Kriefall et al., 2018; Noisette et al., 2016).

### Transcriptomic effects precede detectable phenotypic effects in *C. fornicata* larvae

Gene expression patterns of *C. fornicata* larvae under OA after 4- and 8-d suggested downregulation of genes associated with growth and metabolism and these patterns preceded reductions in larval shell length that became evident at 11- and 12-d (Figure 1A). At 4-d, we observed downregulation of genes associated with mitochondria, ribosomal structures, and electron transport (Figures 4 & S3), which are pathways consistent with reduced growth and reduced respiration (De Wit, Dupont, & Thor, 2016). We also observed downregulation of genes associated with oxidation-reduction processes (Figures 4 & S3), which, in conjunction with downregulation of electron transport, may reflect a dampened immune system. The electron transport chain is a site that produces reactive oxygen species, which are an important internal defense mechanism, but which need to be controlled by antioxidant enzymes (Cao et al., 2018; Liao et al., 2018). Downregulation of genes associated with oxidation-reduction may deplete the availability of antioxidants to counteract reactive oxygen species, thereby overwhelming the antioxidant defense mechanisms. This downregulation could lead to oxidative stress and damages, such as DNA damage and enzyme inactivation, which consequently may impair immune function. At 8-d, larval gene expression changes were not as pronounced, but reductions in ribosomal categories associated with growth were maintained. Despite these GO enrichments at 4- and 8-d, a significant overall transcriptomic effect of pH treatment was not detected (*P* = 0.535; Figure S2B). In addition, the changes in expression of genes associated with growth at 4- and 8-d did not manifest phenotypically until 11-d (Figure 1). Indeed, major gene expression changes have previously been found to precede phenotypic effects in corals under reduced pH (7.6-7.7) over the course of a 28-day experiment (Kaniewska et al., 2012). Overall, our results suggest that the expression of genes associated with metabolism and growth is negatively influenced by OA and that this pattern of gene expression may be a powerful predictor of downstream phenotypic changes.

### Rearing at reduced pH delayed competence for metamorphosis

On days 11 and 12, lower proportions of larvae reared at pH 7.5 and 7.6 metamorphosed within 6 h of induction relative to larvae reared at control conditions (Figure 2), suggesting that OA delays the onset of metamorphic competence in *C. fornicata*. This result recapitulates findings from Kriefall et al. (2018) and Bogan et al. (2019), which found that larval *C. fornicata* exposed to the same reduced pH conditions exhibited significantly longer times to competence for metamorphosis. Delayed onset of metamorphic competence in response to reduced pH has been observed in a wide array of calcifying marine invertebrates, including *Mercenaria mercenaria* (hard clams), *Argopecten irradians* (bay scallops), and *Crassostrea virginica* (Eastern oysters) (Talmage & Gobler, 2009). More importantly, however, competent *C. fornicata* larvae exposed to reduced pH (7.5) were not inhibited from metamorphosis in response to a natural adult-derived cue (Pechenik et al., 2019). Thus, while reduced pH conditions may delay the onset of competence to metamorphose, they do not inhibit already competent larvae from undergoing metamorphosis, further exemplifying *C. fornicata’s* resilience to OA. Given that reduced pH delays the onset of competence, future work might assess whether reduced pH also impacts the length of time required to respond to cues for metamorphosis. Lengthening the time required for a response, like delaying the onset of competence to metamorphose, could increase the potential for predation by lengthening time spent in the water column and increasing risk of mortality (Ross et al., 2016). This additional time in the pelagic environment may also influence a species’ dispersal capacity and potentially lead to range expansion and invasion events, which have been widely observed in *C. fornicata* (Blanchard, 2009; Bohn, Richardson, & Jenkins, 2012).

### Juvenile *C. fornicata* exhibit no carryover effects on growth from reduced pH experienced as larvae

The effects of OA have been shown to frequently transcend life history stages, with exposure at earlier stages influencing development and physiology at later stages (Bogan et al., 2019; Hettinger et al., 2013; Ross et al., 2016). Here, we observed smaller shell lengths for 1-DPM and 4-DPM juveniles that were reared as larvae under reduced pH (Figure S1B-C). However, these reduced shell lengths were a direct consequence of reduced growth observed in larvae at 11- and 12-d (Figure S1A) and no significant difference in rates of juvenile shell growth were observed, despite differences in larval rearing conditions (Figure 3A). Thus, in this experiment, larval pH treatment did not result in carryover effects on growth of juvenile *C. fornicata* that had been transferred to control conditions following metamorphosis. This result contrasts with those of Bogan et al. (2019), who found negative effects of similar larval pH treatments on growth of juvenile *C. fornicata* in three experiments conducted on two different populations across two different seasons. However, those carryover effects were sometimes contingent on co-occurring nutritional deprivation. Similarly, Pechenik and Tyrell (2015) found variation between broods in the carryover effects in juvenile growth resulting from larval nutrition quality. Despite such evidence for intraspecific variation in carryover effects in *C. fornicata*, other studies investigating carryover effects on growth in other molluscs have shown that these effects can persist for months post settlement. For example, Gobler & Talmage (2013) showed that juvenile *Argopecten irradians* (bay scallops) reared at elevated *p*CO_2_ (750 μatm) as larvae experienced reduced growth relative to juveniles reared at ambient *p*CO_2_ (390 μatm) as larvae and these carryover effects did not dissipate for 10 months. Given the intraspecific variation in the expression of carryover effects in *C. fornicata*, future studies should assess the transcriptomic profiles of broods expressing carryover effects and compare them to those of broods not expressing these effects in response to these same conditions.

### Time since pH treatment allowed for recovery of gene expression in juvenile *Crepidula fornicata*

Juvenile transcriptomic analyses suggest that snails exposed to reduced pH as larvae are transiently impacted by OA, but with sufficient time, these effects become negligible (Figures 4 & S3). Juveniles at 1-DPM upregulated genes associated with mitochondria, ribosomal structures, and electron transport, which suggest increased growth, respiration, and metabolism, which are potential carryover effects from conditions experienced as larvae (i.e. juveniles are having to work harder to compensate for prior conditions). It has been suggested that some organisms have the ability to increase rates of their biological processes, including metabolism and growth, to compensate for increased seawater acidity (Gutowska, Melzner, Portner, & Meier, 2010; Lannig, Eilers, Portner, Sokolova, & Bock, 2010; Wood, Spicer, & Widdicombe, 2008). This adaptation, however, comes at a cost and was found to be coupled with reductions in muscle mass in brittlestars (Wood et al., 2008) and a reduction in the incorporation of chitin in the cuttlebones of cephalopods (Gutowska et al., 2010). While we did not observe reduced growth rates in juvenile *C. fornicata* here, it is possible that increased growth and metabolism suggested by the gene expression profiles of juvenile *C. fornicata* similarly involved a phenotypic trade-off that was not quantified in the current study.

In contrast to juveniles at 1-DPM, juveniles at 4-DPM exhibited downregulation of genes associated with mitochondria, ribosomal structures, and oxidation-reduction processes, suggesting reduced growth and respiration. After upregulating genes at 1-DPM in response to pH conditions experienced as larvae, juveniles may downregulate these core pathways at 4-DPM in the absence of pH treatment in order to return to baseline expression levels. Upregulation of genes associated with ciliary structures at 4-DPM suggest that juveniles may be increasing feeding activity upon recovery from pH conditions. Feeding rate is an important determinant of growth and survival, therefore it would be an informative phenotype to quantify in future OA studies on *C. fornicata*. Given that we observed gene expression responses suggestive of dampened immunity in both larvae and juveniles, future studies could assess whether reduced pH results in greater susceptibility to disease and whether prolonged exposure increases susceptibility. Overall, our findings suggest that within a period of 4 days, juvenile *C. fornicata* were able to recover from OA conditions experienced as larvae through acclimation and gene expression plasticity.

### *Crepidula fornicata* larvae exhibited transcriptomic resilience in response to acute pH treatment

Gene expression changes in response to environmental change can occur quickly (Evans & Hofmann, 2012) and monitoring these changes through time can allow for a more in-depth characterization of how stressors influence an organism across short time scales. To test the effects of acute reduced pH conditions, we assessed gene expression responses of *C. fornicata* larvae reared at reduced pH (7.5) relative to larvae reared at higher pH (8.0) over the course of 48 hours. Larval transcriptomes responded rapidly to OA within the first 24-h, upregulating genes associated with mitochondria, ribosomal structures, and oxidation-reduction, suggestive of increased respiration, growth, and metabolism (Figure 5). Regulation of these gene ontology terms have been consistently observed in the literature (reviewed in Strader, Wong, & Hofmann, 2020). For example, Pan, Applebaum, & Manahan (2015) found that *Strongylocentrotus purpuratus* urchin larvae under acidification (~800 μatm *p*CO_2_) increased protein synthesis by approximately 50%. They determined that the majority of available ATP (84%) was accounted for in protein synthesis and ion transport alone, which is indicative of metabolic stress as most energy is directed towards making proteins and offsetting the effects of OA. Basal metabolism was similarly found to increase in the bivalve *Laternula elliptica* after exposure to reduced pH conditions (pH 7.78) over the course of 120 days (Cummings et al., 2011). Our results suggest that increases in expression of genes associated with metabolism and protein synthesis in *C. fornicata* larvae begin to occur at the onset of exposure to reduced pH and that this rapid response may play a role in the ability of the species to acclimatize quickly to OA.

Indeed, we observed reduced overall transcriptomic responses and downregulation of the same terms that were upregulated within the first 24-h by 48-h, which indicates that larvae were able to acclimatize and recover from acute OA treatment (Figure S2). Although physiological measurements were not taken for these larvae, if reared for a longer duration, it is likely that larvae reared in reduced pH (7.5) would have exhibited reduced growth, consistent with the 16-d experiment. Given that no previous study has assessed the transcriptomic effects of OA at hourly scales, these findings highlight the importance of the ephemeral transcriptional changes involved in acclimation to OA that may be overlooked in longer term experiments with coarser sampling resolution.

### *Crepidula fornicata* exhibit resilience to reduced pH conditions through transcriptome plasticity

Overall, our findings suggest that *C. fornicata* exhibit dynamic phenotypic and transcriptomic responses across both larval and juvenile life history stages in response to OA. At the transcriptome level, larvae appeared to experience stress when initially reared in reduced pH treatments (10-h), but almost fully recover baseline transcriptome profiles by 48-h (Figure 5). Similarly, juveniles that were reared under OA conditions as larvae exhibit gene expression patterns consistent with stress at 1-DPM. However, these carryover effects from the larval stage dissipate by 4-DPM (Figures 4 & S3). The ability of *C. fornicata* larvae and juveniles to acclimate to OA quickly sheds light onto the mechanisms underlying this species’ resilience, which likely results from its life spent in variable pH intertidal environments. Reduced larval growth at 11- and 12-d, which appears to be preceded by transcriptomic effects at 4- and 8-d, is further evidence of OA acclimatization. Reduced growth and metabolism may lessen the energetic burden under OA exposure and may provide clues into how these snails are able to exhibit such broad resilience. Overall, our gene expression results in *C. fornicata* corroborate the findings of a recent review by Strader, Wong, & Hofmann (2020), which consistently observed regulation of metabolic processes, calcification and stress response were regulated by marine metazoans in response to OA. In order to fully characterize the dynamic effects of OA on *C. fornicata*, further studies are needed to discern the influence of multiple stressors along with daily and seasonal environmental fluctuations across life history stages (e.g. Rivest, Chen, Fan, Li, & Hofmann, 2017; Waldbusser & Salisbury, 2014). Given that differences in growth and metabolic rates have been found for organisms exposed to static versus fluctuating pH treatments (Britton, Cornwall, Revill, Hurd, & Johnson, 2016; Cornwall et al., 2013; Mangan, Urbina, Findlay, Wilson, & Lewis, 2017; Roleda et al., 2015), we suggest that future work should examine the phenotypic and transcriptomic responses across different treatment durations and variable/static OA conditions.

## Acknowledgements

We thank C. Gillespie, M. Lee, J. Litle, and J. Trudel for assisting with maintenance of larval cultures and R. Guenther for support of seawater conditioning and chemical analysis in the Ocean Acidification Environmental Laboratory at the University of Washington Friday Harbor Laboratories. D. Cooper at Taylor Shellfish Co. kindly provided access to our adult field collection site. We thank N. Kriefall for insightful discussions about the biological meaning of the study and F. Giler for assisting with development of figures.

## Author Contributions

AP, JAP and SWD designed the experiment. ML, AP and JAP conducted the experiment. BEB and CLR completed all molecular work and TagSeq library preparations. CLR and XC performed all statistical and bioinformatic analyses with supervision from SWD. CLR and SWD drafted the manuscript. All authors edited and approved the manuscript.

## Data Accessibility Statement

Detailed protocol of library preparation and bioinformatics can be found at https://github.com/z0on/tag-based_RNAseq. Protocol for gene ontology analysis can be found at https://github.com/z0on/GO_MWU. Annotation files for the *Crepidula fornicata* transcriptome are available at http://sites.bu.edu/davieslab/data-code. Raw reads have been submitted to SRA under PRJNA549522. All files from the gene expression analyses and all phenotypic measurement datasets are available at https://github.com/chrislreyes/Crepidula.

