## Supplemental Information for "The marine gastropod *Crepidula fornicata* remains resilient to ocean acidification across two life history stages"

Figure S1 | Mean larval shell length (µm) in each pH treatment measured at 4, 8, 11, and 12 days in treatment. Mean juvenile shell length growth rates (µm/day) measured 24 hours post settlement (1-DPM) and 4 days later (4-DPM). Error bars represent +/- one standard error and different letters for pH treatments indicate significantly different means based on Tukey's HSD tests ($P < 0.05$).

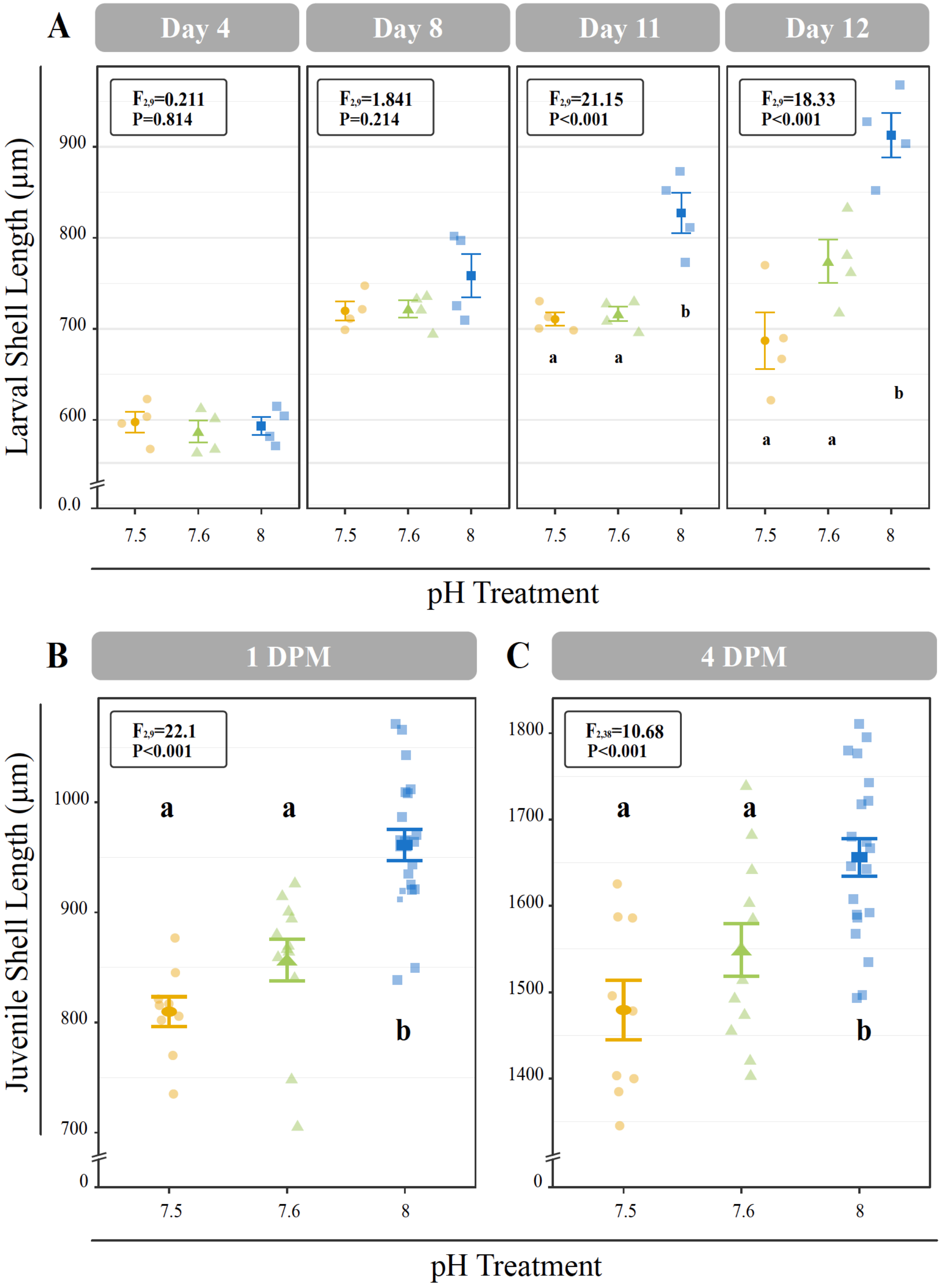

Figure S2 | Canonical Correspondence Analysis (CCA) of all log transformed isogroups clustered by time point for larvae in the 48-h experiment (A) and 16-d experiment (B) and clustered by treatment for juveniles in the 16-d experiment (C). Time point was found to be more impactful on larvae and treatment more impactful on juveniles, but overall, responses of C. *fornicata* larvae and juveniles across time points and pH treatments were found to not be significant. For larvae (A-B), shapes indicate time point (4-, 10-, 24-, and 48- h (A) and 4- and 8- d (B)) and colors indicate pH treatment condition: blue = pH 8.0, green = pH 7.6, yellow = pH 7.5. For juveniles (C), shapes indicate pH treatment (7.5, 7.6, 8.0) and colors indicate time point: blue = 1-DPM, purple = 4-DPM.

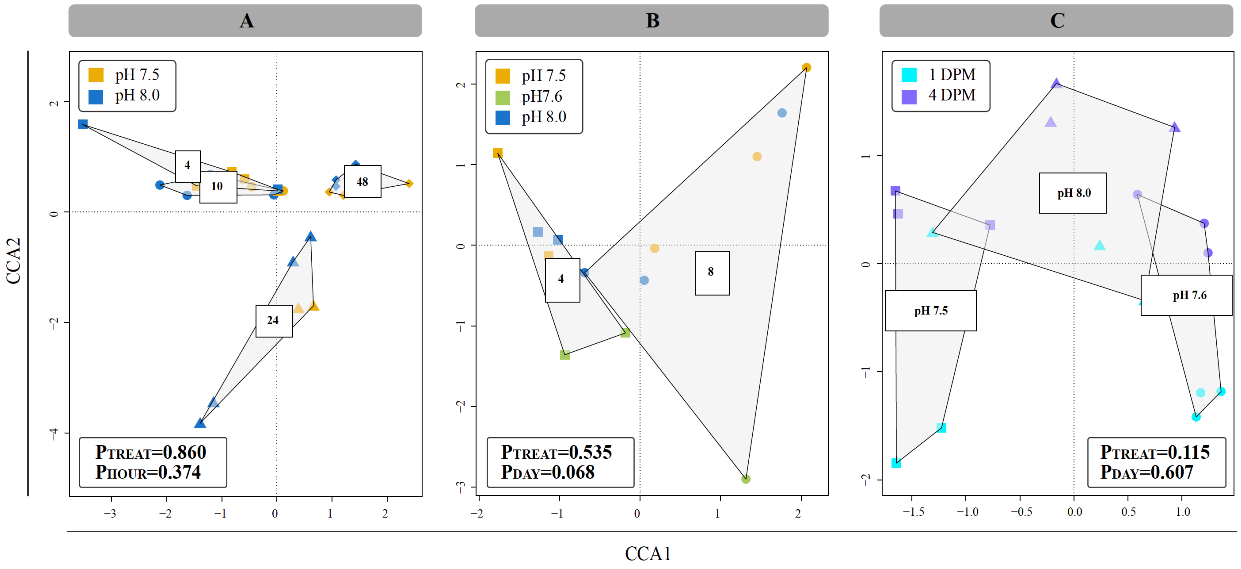

Figure S3 | Significantly enriched gene ontology (GO) categories for the pairwise comparison between pH 7.6 and pH 8.0 treatments for larvae and juveniles in the 16-d experiment. Mann-Whitney U (MWU) tests were conducted based on ranking of signed log p-values and the results were plotted as dendograms with an indication of genes shared between categories. Enrichment by 'cellular component', 'biological process', and 'molecular function' are shown for 4- and 8-d for larvae and for 1-DPM and 4-DPM for juveniles. Overrepresented categories relative to pH 8.0 are colored as red and underrepresented categories are colored as blue. A blank grid indicates that there were no significantly enriched categories for that division at that time point.

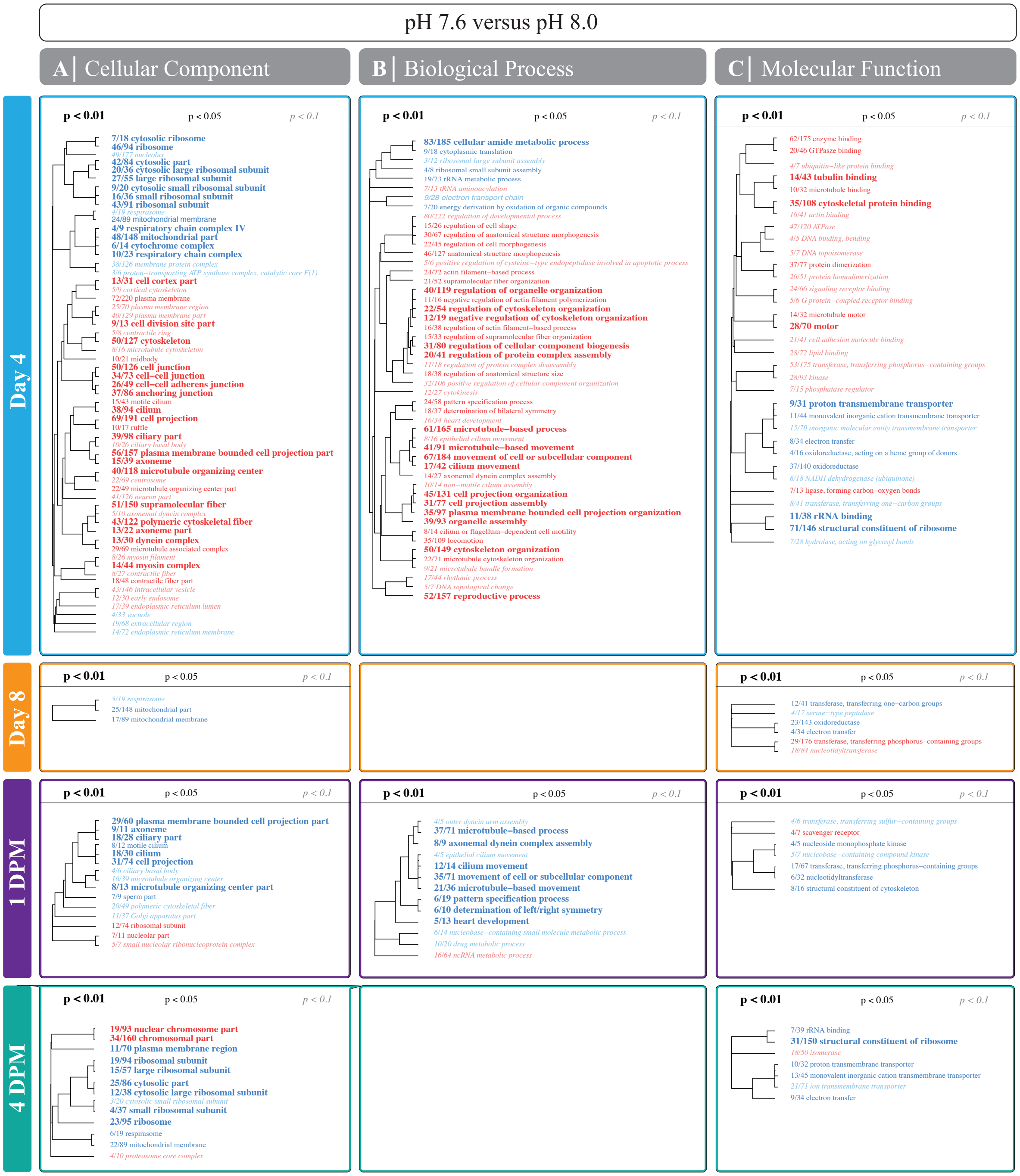

pH 7.6 versus pH 8.0
A | Cellular Component
B | Biological Process
C | Molecular Function
Day 4
p < 0.01
p < 0.05
p < 0.1
7/18 cytosolic ribosome
46/94 ribosome
49/177 nucleolus
42/84 cytosolic part
20/36 cytosolic large ribosomal subunit
27/55 large ribosomal subunit
9/20 cytosolic small ribosomal subunit
16/36 small ribosomal subunit
43/91 ribosomal subunit
4/19 respirasome
24/89 mitochondrial membrane
4/9 respiratory chain complex IV
48/148 mitochondrial part
6/14 cytochrome complex
10/23 respiratory chain complex
38/126 membrane protein complex
3/6 proton−transporting ATP synthase complex, catalytic core F(1)
13/31 cell cortex part
5/9 cortical cytoskeleton
72/220 plasma membrane
25/70 plasma membrane region
40/129 plasma membrane part
9/13 cell division site part
5/8 contractile ring
50/127 cytoskeleton
8/16 microtubule cytoskeleton
10/21 midbody
50/126 cell junction
34/73 cell−cell junction
26/49 cell−cell adherens junction
37/86 anchoring junction
15/43 motile cilium
38/94 cilium
69/191 cell projection
10/17 ruffle
39/98 ciliary part
10/26 ciliary basal body
56/157 plasma membrane bounded cell projection part
15/39 axoneme
40/118 microtubule organizing center
22/69 centrosome
22/49 microtubule organizing center part
41/126 neuron part
51/150 supramolecular fiber
5/10 axonemal dynein complex
43/122 polymeric cytoskeletal fiber
13/22 axoneme part
13/30 dynein complex
29/69 microtubule associated complex
8/26 myosin filament
14/44 myosin complex
8/27 contractile fiber
18/48 contractile fiber part
43/146 intracellular vesicle
12/30 early endosome
17/39 endoplasmic reticulum lumen
4/33 vacuole
19/68 extracellular region
14/72 endoplasmic reticulum membrane
83/185 cellular amide metabolic process
9/18 cytoplasmic translation
3/12 ribosomal large subunit assembly
4/8 ribosomal small subunit assembly
19/73 rRNA metabolic process
7/13 tRNA aminoacylation
8/28 electron transport chain
7/20 energy derivation by oxidation of organic compounds
80/222 regulation of developmental process
15/26 regulation of cell shape
30/67 regulation of anatomical structure morphogenesis
22/45 regulation of cell morphogenesis
46/127 anatomical structure morphogenesis
5/6 positive regulation of cysteine−type endopeptidase involved in apoptotic process
24/72 actin filament−based process
21/52 supramolecular fiber organization
40/119 regulation of organelle organization
11/16 negative regulation of actin filament polymerization
22/54 regulation of cytoskeleton organization
12/19 negative regulation of cytoskeleton organization
16/38 regulation of actin filament−based process
15/33 regulation of supramolecular fiber organization
31/80 regulation of cellular component biogenesis
20/41 regulation of protein complex assembly
11/18 regulation of protein complex disassembly
18/38 regulation of anatomical structure size
32/106 positive regulation of cellular component organization
12/27 cytokinesis
24/58 pattern specification process
18/37 determination of bilateral symmetry
16/34 heart development
61/165 microtubule−based process
8/16 epithelial cilium movement
41/91 microtubule−based movement
67/184 movement of cell or subcellular component
17/42 cilium movement
14/27 axonemal dynein complex assembly
10/14 non−motile cilium assembly
45/131 cell projection organization
31/77 cell projection assembly
35/97 plasma membrane bounded cell projection organization
39/93 organelle assembly
8/14 cilium or flagellum−dependent cell motility
35/109 locomotion
50/149 cytoskeleton organization
22/71 microtubule cytoskeleton organization
9/21 microtubule bundle formation
17/44 rhythmic process
5/7 DNA topological change
52/157 reproductive process
62/175 enzyme binding
20/46 GTPase binding
4/7 ubiquitin−like protein binding
14/43 tubulin binding
10/32 microtubule binding
35/108 cytoskeletal protein binding
16/41 actin binding
47/120 ATPase
4/5 DNA binding, bending
5/7 DNA topoisomerase
37/77 protein dimerization
26/51 protein homodimerization
24/66 signaling receptor binding
5/6 G protein−coupled receptor binding
14/32 microtubule motor
28/70 motor
21/41 cell adhesion molecule binding
28/72 lipid binding
53/175 transferase, transferring phosphorus−containing groups
28/93 kinase
7/15 phosphatase regulator
9/31 proton transmembrane transporter
11/44 monovalent inorganic cation transmembrane transporter
15/70 inorganic molecular entity transmembrane transporter
8/34 electron transfer
4/16 oxidoreductase, acting on a heme group of donors
37/140 oxidoreductase
6/18 NADH dehydrogenase (ubiquinone)
7/13 ligase, forming carbon−oxygen bonds
8/41 transferase, transferring one−carbon groups
11/38 rRNA binding
71/146 structural constituent of ribosome
7/28 hydrolase, acting on glycosyl bonds
Day 8
5/19 respirasome
25/148 mitochondrial part
17/89 mitochondrial membrane
12/41 transferase, transferring one−carbon groups
4/17 serine−type peptidase
23/143 oxidoreductase
4/34 electron transfer
29/176 transferase, transferring phosphorus−containing groups
18/84 nucleotidyltransferase
1 DPM
29/60 plasma membrane bounded cell projection part
9/11 axoneme
18/28 ciliary part
8/12 motile cilium
18/30 cilium
31/74 cell projection
4/6 ciliary basal body
16/39 microtubule organizing center
8/13 microtubule organizing center part
7/9 sperm part
20/49 polymeric cytoskeletal fiber
11/37 Golgi apparatus part
12/74 ribosomal subunit
7/11 nucleolar part
5/7 small nucleolar ribonucleoprotein complex
4/5 outer dentin arm assembly
37/71 microtubule−based process
8/9 axonemal dynein complex assembly
4/5 epithelial cilium movement
12/14 cilium movement
35/71 movement of cell or subcellular component
21/36 microtubule−based movement
6/19 pattern specification process
6/10 determination of left/right symmetry
5/13 heart development
6/14 nucleobase−containing small molecule metabolic process
10/20 drug metabolic process
16/64 ncRNA metabolic process
4/6 transferase, transferring sulfur−containing groups
4/7 scavenger receptor
4/5 nucleoside monophosphate kinase
5/7 nucleobase−containing compound kinase
17/67 transferase, transferring phosphorus−containing groups
6/32 nucleotidyltransferase
8/16 structural constituent of cytoskeleton
4 DPM
19/93 nuclear chromosome part
34/160 chromosomal part
11/70 plasma membrane region
19/94 ribosomal subunit
15/57 large ribosomal subunit
25/86 cytosolic part
12/38 cytosolic large ribosomal subunit
3/20 cytosolic small ribosomal subunit
4/37 small ribosomal subunit
23/95 ribosome
6/19 respirasome
22/89 mitochondrial membrane
4/10 proteasome core complex
7/39 rRNA binding
31/150 structural constituent of ribosome
18/50 isomerase
10/32 proton transmembrane transporter
13/45 monovalent inorganic cation transmembrane transporter
21/71 ion transmembrane transporter
9/34 electron transfer

**Supplemental Tables**

Table S1 | Characteristics of seawater used for larval culturing (12 days) during the 16 d experiment. Values of pH are reported (from left) as the nominal treatment target values, the actual values for new seawater added to cultures, and the values recorded immediately before regular seawater changes. TA, total alkalinity; ΩAr, saturation state of aragonite; pCO2, partial pressure of carbon dioxide.

| Target pH | Value | pH (total) | Salinity (ppt) | Temp. (°C) | TA (µmol/kg) | $\Omega_{Ar}$ | $pCO_2$ (µatm) | pH at change (total) |
|---|---|---|---|---|---|---|---|---|
| 8.0 | Mean | 8.00 | 28.92 | 20.12 | 2071.3 | 2.14 | 436.1 | 7.93 |
| | SD | 0.01 | 0.60 | 0.11 | 26.5 | 0.06 | 16.1 | 0.03 |
| | n | 6 | 6 | 6 | 4 | 4 | 4 | 6 |
| 7.6 | Mean | 7.59 | 29.03 | 20.16 | 2071.3 | 0.94 | 1217.1 | 7.52 |
| | SD | 0.01 | 0.65 | 0.08 | 26.5 | 0.04 | 25.2 | 0.05 |
| | n | 6 | 6 | 6 | 4 | 4 | 4 | 6 |
| 7.5 | Mean | 7.53 | 29.05 | 20.20 | 2071.3 | 0.81 | 1444.7 | 7.48 |
| | SD | 0.01 | 0.69 | 0.14 | 26.5 | 0.03 | 37.4 | 0.03 |
| | n | 6 | 6 | 6 | 4 | 4 | 4 | 6 |

Table S2 | Characteristics of seawater used during the 48 h experiment. Values of pH are reported (from left) as the nominal treatment target values, and the actual values for new seawater added to cultures at the beginning of the experiment (0 h) or measured in 4 replicate cultures after 4, 10, 24, and 48h. TA, total alkalinity; $\Omega_{Ar}$, saturation state of aragonite; $pCO_2$, partial pressure of carbon dioxide.

| Target pH | time (h) | Value | pH (total) | Salinity (ppt) | Temp. (°C) | TA (µmol/kg) | $\Omega_{Ar}$ | $pCO_2$ (µatm) |
|---|---|---|---|---|---|---|---|---|
| 8.0 | 0 | Initial | 8.05 | 29.6 | 20.1 | 2101.7 | 2.45 | 378.6 |
| | 4 | Mean | 8.06 | 29.65 | 20.38 | | | |
| | | SD | 0.01 | 0.06 | 0.15 | | | |
| | 10 | Mean | 8.06 | 29.65 | 20.23 | | | |
| | | SD | 0.01 | 0.06 | 0.10 | | | |
| | 24 | Mean | 8.02 | 29.40 | 20.25 | | | |
| | | SD | 0.02 | 0.41 | 0.34 | | | |
| | 48 | Mean | 8.01 | 29.50 | 19.50 | | | |
| | | SD | 0.01 | 0.16 | 0.08 | | | |
| 7.5 | 0 | Initial | 7.54 | 29.6 | 19.9 | 2101.7 | 0.86 | 1398.3 |
| | 4 | Mean | 7.57 | 29.63 | 20.23 | | | |
| | | SD | 0.02 | 0.13 | 0.10 | | | |
| | 10 | Mean | 7.57 | 29.63 | 20.05 | | | |
| | | SD | 0.02 | 0.05 | 0.06 | | | |
| | 24 | Mean | 7.52 | 29.68 | 19.98 | | | |
| | | SD | 0.02 | 0.10 | 0.10 | | | |
| | 48 | Mean | 7.55 | 29.43 | 19.50 | | | |
| | | SD | 0.02 | 0.13 | 0.08 | | | |

Table S3 | Summary of RNA libraries for the 16-d experiment, including raw single-end reads, trimmed reads, mapped counts, and mapping efficiencies (%). Samples found to be outliers during gene expression analyses and excluded from subsequent analyses are colored red.

| Day | Sample | Raw Reads | Trimmed Reads | Total Counts | % Mapped |
|---|---|---|---|---|---|
| 4 | 7.5A | 20031800 | 2715462 | 1142969 | 42.1 |
| 4 | 7.5B | 11301574 | 1477813 | 665524 | 45.0 |
| 4 | 7.6B | 17231518 | 3107244 | 1327316 | 42.7 |
| 4 | 7.6C | 12365225 | 2108339 | 932421 | 44.2 |
| 4 | 8.0B | 13650429 | 2171171 | 954244 | 44.0 |
| 4 | 8.0C | 20935860 | 3350460 | 1468909 | 43.8 |
| 8 | 7.5B | 8655806 | 2362028 | 1003391 | 42.5 |
| 8 | 7.5C | 17879591 | 3135665 | 1368268 | 43.6 |
| 8 | 7.5D | 8148648 | 2083059 | 918446 | 44.1 |
| 8 | 7.6B | 2345757 | 755538 | 330871 | 43.8 |
| **8** | **7.6C** | **2789212** | **428839** | **199558** | **46.5** |
| **8** | **7.6D** | **490776** | **8980** | **4380** | **48.8** |
| 8 | 8.0A | 13214356 | 2092572 | 902746 | 43.1 |
| 8 | 8.0C | 13232434 | 2450088 | 1055424 | 43.1 |
| 8 | 8.0D | 13696321 | 2497771 | 1063168 | 42.6 |
| 1DPM | 7.5A | 13917737 | 2196945 | 972423 | 44.3 |
| 1DPM | 7.5B | 10736059 | 1738802 | 750239 | 43.1 |
| **1DPM** | **7.5C** | **620625** | **8475** | **4176** | **49.3** |
| 1DPM | 7.6A | 9456989 | 2735258 | 1156530 | 42.3 |
| 1DPM | 7.6B | 8785582 | 2383939 | 1008800 | 42.3 |
| **1DPM** | **7.6C** | **986675** | **265572** | **125367** | **47.2** |
| 1DPM | 7.6D | 6713990 | 2124722 | 905133 | 42.6 |
| 1DPM | 8.0A | 12604899 | 3440782 | 1404888 | 40.8 |
| 1DPM | 8.0B | 13689368 | 2748730 | 1154117 | 42.0 |
| 1DPM | 8.0D | 8562743 | 2107538 | 887547 | 42.1 |
| 4DPM | 7.5B | 5187391 | 1703006 | 733385 | 43.1 |
| 4DPM | 7.5C | 18056592 | 3395007 | 1459340 | 43.0 |
| 4DPM | 7.5D | 15190058 | 2949961 | 1255646 | 42.6 |
| 4DPM | 7.6A | 10159287 | 3192001 | 1299882 | 40.7 |
| 4DPM | 7.6C | 9303499 | 2718200 | 1146724 | 42.2 |
| 4DPM | 7.6D | 12933223 | 3781562 | 1557855 | 41.2 |
| **4DPM** | **8.0A** | **14494** | **3855** | **1829** | **47.4** |
| 4DPM | 8.0B | 7939351 | 2523340 | 1070388 | 42.4 |
| 4DPM | 8.0C | 7451760 | 2423257 | 987887 | 40.8 |
| 4DPM | 8.0D | 13088795 | 3411320 | 1381910 | 40.5 |

Table S4 | Summary of RNA libraries for the 48-h experiment, including raw single-end reads, trimmed reads, mapped counts, and mapping efficiencies (%). Samples found to be outliers during gene expression analyses and excluded from subsequent analyses are colored red.

| **Hour** | **Sample** | **Raw Reads** | **Trimmed Reads** | **Total Counts** | **% Mapped** |
|---|---|---|---|---|---|
| 0 | 8.0A | 3398321 | 908567 | 402011 | 44.2 |
| 0 | 8.0B | 3874971 | 981793 | 437684 | 44.6 |
| 0 | 8.0C | 4077075 | 888164 | 407762 | 45.9 |
| 0 | 8.0D | 1103488 | 298350 | 126365 | 42.4 |
| 4 | 7.5A | 10758493 | 1430476 | 605458 | 42.3 |
| 4 | 7.5B | 9913935 | 2087837 | 916263 | 43.9 |
| 4 | 7.5C | 5944596 | 733095 | 314277 | 42.9 |
| 4 | 7.5D | 5287317 | 724507 | 324889 | 44.8 |
| 4 | 8.0A | 6539817 | 2147901 | 926678 | 43.1 |
| **4** | **8.0B** | **956622** | **98985** | **36646** | **37.0** |
| 4 | 8.0C | 4901613 | 203310 | 94004 | 46.2 |
| **4** | **8.0D** | **1008450** | **46607** | **16751** | **35.9** |
| 10 | 7.5A | 8118936 | 903449 | 377236 | 41.8 |
| 10 | 7.5B | 4652645 | 1151415 | 522103 | 45.3 |
| 10 | 7.5C | 4751702 | 1442681 | 621347 | 43.1 |
| **10** | **7.5D** | **656692** | **65948** | **21902** | **33.2** |
| 10 | 8.0A | 7555520 | 501985 | 223739 | 44.6 |
| 10 | 8.0B | 4028652 | 536405 | 225729 | 42.1 |
| 10 | 8.0C | 4839052 | 626000 | 260243 | 41.6 |
| 10 | 8.0D | 5104318 | 1368157 | 587962 | 43.0 |
| 24 | 7.5A | 1825175 | 677711 | 306236 | 45.2 |
| **24** | **7.5B** | **901923** | **105238** | **31495** | **29.9** |
| 24 | 7.5C | 2841801 | 802780 | 360527 | 44.9 |
| **24** | **7.5D** | **1374012** | **149658** | **48236** | **32.2** |
| 24 | 8.0A | 3925847 | 432207 | 178560 | 41.3 |
| 24 | 8.0B | 3604435 | 1260590 | 554924 | 44.0 |
| 24 | 8.0C | 2518079 | 337301 | 139452 | 41.3 |
| 24 | 8.0D | 8629343 | 2027985 | 891746 | 44.0 |
| 48 | 7.5A | 12084816 | 1498925 | 609382 | 40.7 |
| 48 | 7.5B | 10460064 | 2874416 | 1235041 | 43.0 |
| 48 | 7.5C | 1517676 | 431402 | 178275 | 41.3 |
| 48 | 7.5D | 2569918 | 680018 | 297157 | 43.7 |
| 48 | 8.0A | 3683716 | 1008282 | 443670 | 44.0 |
| 48 | 8.0B | 9576664 | 2118374 | 917558 | 43.3 |
| 48 | 8.0C | 4334377 | 696262 | 289920 | 41.6 |
| 48 | 8.0D | 5150192 | 1408095 | 620377 | 44.1 |
